# Transcriptomic regulation of juvenile-to-adult vegetative phase transition in grapevine

**DOI:** 10.1101/2024.12.29.630313

**Authors:** Yolanda Ferradás, Carolina Royo, José Miguel Martínez-Zapater, Diego Lijavetzky

**Affiliations:** Departamento de Biología Funcional, Universidade de Santiago de Compostela, Santiago de Compostela, Spain; Instituto de Ciencias de la Vid y del Vino, ICVV, CSIC - Universidad de La Rioja - Gobierno de La Rioja, Logroño 26007, La Rioja, Spain; Instituto de Biología Agrícola de Mendoza (IBAM, CONICET-UNCUYO), Almirante Brown 500, M5528AHB. Chacras de Coria, Mendoza, Argentina

**Author notes:** Both authors contributed equally to the work (Yolanda Ferradás, Carolina Royo).

**Keywords:** Grapevine, Juvenile-to-adult phase change, Flowering transition, Tendril development, miRNA, Transcriptomic analysis

## Abstract

Plants go through two distinct stages in their vegetative phase, with the juvenile stage being characterized by a lack of maturity to respond to flowering induction stimuli and the adult stage marked by the presence of this capacity. Phase transition has been extensively analysed in herbaceous species such as Arabidopsis and maize, where the sequential activity of miR156 and miR172 in the control of the juvenile to adult phase transition has been determined. Contrarily, little is known about most woody perennial crops, where phase transition appears to be dissociated, with a first transition from juvenile to adult vegetative state in the first year and a subsequent induction to flower in later years under flowering-inductive environmental conditions. This significantly extended vegetative phase makes fruit production depend on the grafting of adult vegetative materials. A particular aspect of grapevine vegetative phase transition is that it is marked by the differentiation of tendrils, a modified sterile reproductive organ adapted to climbing, which is continuously generated with different patterns in different *Vitis* species. When the grapevine plant reaches flowering inductive condition in later years, it produces inflorescences in place of some tendrils. As a first step to understand the regulation of phase change in grapevine, we have performed a detailed gene expression analysis of the juvenile-to-adult phase transition during the development of grapevine plantlets grown from seeds. The RNA-seq analysis demonstrated that miR156 was significantly repressed in the grapevine’s adult phase, where the appearance of tendrils acts as a marker of the transition. Consistent with the results reported in other species, we observed the activation of several *SPL* genes, known to be targets of miR156, providing evidence for the conservation of the miR156-SPLs regulatory module in grapevine. However, no variation was detected in the expression of miR172 and *TPS* genes were found downregulated, two key determinants in the transition to flowering in other species. This could be explained considering that grapevines do not flower during the first years of growth. Interestingly, we were able to observe the overexpression of several genes known to be involved in the establishment of flower meristem identity, which in the case of grape had also been detected along tendril development, consistent with the proposed common ontogenetic origin of tendrils and inflorescences in the *Vitaceae* family.

## Introduction

Developmental genetic programs, which are regulated by both endogenous and exogenous stimuli, govern the transition between the distinct growth phases that comprise the life cycle of flowering plants. After germination and prior to attaining reproductive maturity, most plant shoots undergo a period of juvenile vegetative growth. As juvenile annual plants move towards reproductive competence also known as adult vegetative phase, they show progressive developmental changes in their vegetative organs such as dimension, shape and phyllotaxis of the leaves, changes in internode elongation, or in the patterns of trichome distribution (Huijser & Schmid, 2011; M. Keller, 2020b). When a well-defined set of environmental and endogenous conditions triggers the initiation of reproductive development, such as photoperiod, vernalization or age, the adult vegetative plants initiate the development of flowers (Poethig, 1990, 2010). It remains uncertain if the same variables regulating the juvenile-to-adult phase transition in annual plant species also regulate this transition in woody perennial plants. Shrubs and trees frequently display more pronounced distinctions between their juvenile and adult stages, with woody plants generally revealing these differences more consistently than herbaceous species. Secondly, the juvenile and adult vegetative stages are relatively short in herbaceous plants like *Arabidopsis thaliana* and maize, whereas they can persist for several years in perennial species (Lawson & Poethig, 1995a; Wang et al., 2011). Understanding this regulation is particularly essential in fruit trees, since an extended juvenile phase severely preclude crop selection and improvement, given that breeding and production features depend on attaining reproductive maturity, which includes flowering and fruiting (Ahsan et al., 2019). This significantly extended vegetative phase makes fruit production depend on the grafting of adult vegetative materials.

It has been evident in recent years that there are several key regulatory variables shared by the networks that govern the transitions from the juvenile-to-adult phases. Furthermore, several of these factors influence specific heteroblastic characteristics that differentiate between juvenile and adult phases in annual plants (Poethig, 2010). At the transcriptional level, two highly conserved microRNAs (miR156 and miR172) have been widely identified as essential elements of the genetic control mechanisms underlying phase variations in several plant species like *A. thaliana* (Wu & Poethig, 2006), maize (Chuck et al., 2007), rice, (Xie et al., 2012) and tomato (Silva et al., 2014). The expression of miR156 is highly abundant in seedlings of the model plant *A. thaliana*, and it declines as growth progresses, whereas miR172 exhibits an opposite expression pattern. The inhibition of miR156 during the transition from the juvenile stage to the adult stage enables overexpression of several SQUAMOSA PROMOTER BINDING PROTEIN-LIKE (SPL) transcription factors that are negatively regulated by miR156 (Wu et al., 2009; Wu & Poethig, 2006; Xu, Hu, Zhao, et al., 2016). Since miR172 is directly targeted by *SPL* genes, its expression increases gradually during the transition to the adult phase. In turn, miR172 participates in floral induction by directly repressing the expression of six Arabidopsis AP2-like transcriptional repressors including *APETALA2 (AP2), TARGET OF EARLY ACTIVATION TAGGED (EAT) 1 (TOE1), TOE2, TOE3, SCHLAFMUTZE (SMZ), and SCHNARCHZAPFEN (SNZ)*. These proteins are responsible for the suppression of *SOC1 and FT*, which are activated during floral induction by the increasing the activity of miR172 (Aukerman & Sakai, 2003b; Chen, 2004). Such expression activation of *SOC1* (and various *SPLs*), allows the upregulation of different floral meristem identity genes, such as *APETALA1* (*AP1*), *FRUITFULL* (*FUL*), and *LEAFY* (*LFY*) promoting flower meristem development (Lee et al., 2008; Liu et al., 2008; Xu, Hu, Zhao, et al., 2016). The conservation of the molecular regulation of these transition from the juvenile-to-adult phases in maize, rice and tomato is mediated by the corresponding orthologous genes and gene families (Chuck et al., 2007; Silva et al., 2014; Xie et al., 2012).

Despite the well-known fact that vegetative development is regulated by a temporal decrease in *miR156* levels, how this decreased expression is initiated and then maintained during shoot development remains elusive. Recent research suggests miR159 and MYB33 function as moderators of vegetative phase change. Specifically, miR159 facilitates vegetative phase change by repressing MYB33 expression, targeting *MYB33* transcripts for degradation in Arabidopsis. This prevents MYB33 from hyperactivating *miR156* expression throughout shoot development, ensuring correct timing of the juvenile-to-adult transition (Guo et al., 2017). Sugar metabolism was also associated with the promotion of phase transition through the repression of miR156 expression. In this sense, the repression of miR156 induced by glucose was dependent on the signaling activity of HEXOKINASE1 (HXK1) (Yang et al., 2013). Moreover, carbohydrate status was also involved in flowering induction, where TREHALOSE-6-PHOSPHATE SYNTHASE 1 (TPS1) seems to be required for the timely initiation of flowering (Wahl et al., 2013).

Gibberellins (GAs) are involved in the regulation of plant growth and are also known as adult phase promoters in Arabidopsis and maize (Lawson & Poethig, 1995b; Telfer & Poethig, 1998). The gibberellin pathway, which is required for flowering promotion in Arabidopsis, induces LFY expression, independent of photoperiod (Blazquez & Weigel, 2000). Additionally, by using the rice *d18-dy* mutant, Tanaka (2012) has shown that GAs participate in the change from juvenile to adult phase independent of microRNA regulation, particularly miR156 and miR172.

Knowledge about the genetic regulation of phase transition in perennial species is less abundant than in annuals. However, Wang et al. (2011), working with different woody trees, and Ahsan et al. (2019), working with horticultural tree crops, coincided in remarking that miR156 is an evolutionarily conserved regulator of vegetative phase change. However, in both reports, the role of miR172 was less clear than in annual plants, while the expression of flowering transition regulators was not deeply evaluated (Ahsan et al., 2019; Wang et al., 2011).

When grapevine (*Vitis vinifera* L.) plants are grown from seeds, vegetative development involves a transition from a juvenile to an adult phase after the formation of 6–10 nodes. This phase change is marked by a shift from spiral to alternate phyllotaxis, changes in leaf morphology from round to lobed forms, and the development of lateral meristems, known as uncommitted primordia, which arise opposite to leaf primordia and initially form tendrils. As in other Vitaceae species, grapevine is characterized by a climbing growth habit supported on tendrils, which share a common ontogenetic origin with inflorescences (Lacroix & Posluszny, 1989; Mullins et al., 1992). In later years, under flowering-inductive conditions, some of those lateral uncommitted primordia will produce inflorescences instead of tendrils, depending on the environmental conditions and the plant status, related to the levels of bioactive gibberellins (Boss & Thomas, 2002; Gerrath, 1992; M. Keller, 2020a; Srinivasan & Mullins, 1980). While gene expression related to the flowering transition has been studied in grapevine (Carmona et al., 2008; Díaz-Riquelme et al., 2014), no studies have yet analyzed the vegetative transition from the juvenile to the adult phase.

To understand the transcriptional regulation of the juvenile-to-adult phase transition during the development of grapevine plantlets grown from seeds, we performed a global gene expression analysis of plants derived from the self-cross of the near-homozygous line PN40024 line (Jaillon et al., 2007; Velt et al., 2023), allowing a replicated RNA-analysis of different developmental stages. The quasi-homozygous state of this line warrants that all derived selfed-plantlets share a very similar genetic composition and behave similarly regarding the timing of phase transition.

## Materials and methods

### 1. Plant material

Plants from the PN40024 near-homozygous line (Jaillon et al., 2007; Velt et al., 2023) grown at the Grapevine Germplasm Collection of the Instituto de Ciencias de la Vid y del Vino (ICVV; ESP-217) at Finca La Grajera (Logroño, La Rioja, Spain) were self-pollinated to generate the plant material required for the present work (PN40024_SP plants). Seeds from three bunches were stirring in water to remove the pulp and rinsed with distilled water several times, air-dried for 24 h, treated with CuOH2 and stored at 4°C. Before sowing, seeds were treated with 10% H2SO4, rinsed and stored in a mix 1:1:1 of (soil:peat moss:vermiculite) for 90 days at 4°C with constant humidity. After a warm stratification (30-36°C for 2 days), seeds were germinated in a grow chamber (Aralab, https://aralab.pt/) with 16 h of light at 25°C and 8 h of darkness at 22°C with 80% of constant humidity.

### 2. Sampling, experimental design and RNA isolation

Previous observations indicated that tendril primordia appeared in these self-pollinated PN40024 plants after the 8-10 node. For these reasons, apex from plants with 6-8 nodes were considered as juvenile while apex from plants with 8-10 nodes displaying one detectable tendril primordia were considered to be already in the adult vegetative phase. We collected shoot apexes from juvenile and adult seed-derived plantlets for the RNA-seq analysis. Each of the six biological replicates (three juvenile and three adult) were composed from the shoot apex of three individual plants, using 18 plantlets in total. Samples from the adult plants were collected after the appearance of the first tendril from plants with 8-10 nodes (Supplementary Figure 1). Juvenile samples were collected from plants with 6-8 nodes before appearance of the first tendril (Lacroix & Posluszny, 1989). For the second year RT-qPCR experiments, samples were collected from apex of greenhouse two-year plants, after and before the detection of the first tendril. Samples were frozen in liquid nitrogen and stored at −80°C until use. Total RNA was extracted from frozen apexes using a Spectrum™ Plant Total RNA kit (Sigma-Aldrich). An in-column DNase digestion step with the RNase-Free DNase Set (Qiagen) was added. RNA quantification was performed by means of a Nanodrop 2000 Spectrophotometer (Thermo Fisher Scientific), while integrity and potential genomic DNA contaminations were checked through agarose gel electrophoresis.

### 3. Library construction and RNA sequencing

Six samples (two developmental stages × three biological replicates) were sent to the CRG Genomic Unit (Barcelona, Spain). Once in destination they were checked for total RNA integrity using a Bioanalyzer RNA Nano 6000 chip. All the samples qualified to proceed with the library construction having an RNA Integrity Number (RIN) ≥ 9. NGS transcriptomic libraries were constructed using a TruSeq Stranded mRNA LT Sample Prep Kit (Illumina). To verify the size of PCR enriched fragments, the template size distribution was checked on an Agilent Technologies 2100 Bioanalyzer using a DNA 1000 chip. The sequencing of libraries was performed as paired-end 150 bp reads on an Illumina NextSeq platform.

The base quality of the raw reads was checked with FastQC (Andrews, 2010). Trimming of sequencing adaptors and low quality sequences was performed with Trimmomatic (Bolger et al., 2014) using ‘ILLUMINACLIP:TruSeq3-PE. fa:2:30:10 LEADING:21 TRAILING:21 MINLEN:30’ as parameters. Reads corresponding to rRNA and tRNA were removed from the trimmed Illumina reads using Bowtie2 (Langmead & Salzberg, 2012) aligner, by mapping the reads to a specific grapevine database (Chan & Lowe, 2009). Clean filtered reads were aligned on the grapevine PN40024.v4 genome (Velt et al., 2023), using the STAR aligner (Dobin et al., 2013) with ‘--alignIntronMin 20 --alignIntronMax 20,000 --outSAMtype BAM SortedByCoordinate -- outReadsUnmapped Fastx’ as parameters. Subsequently, the aligned reads were assembled by means of StringTie (Pertea et al., 2015) and new transcripts were extracted and annotated using the GffCompare (Pertea & Pertea, 2020) program (GffCompare classes “u”, “x”, to adjust). The assembly of new transcripts and their classification into coding and noncoding genes were performed according to Chialva et al. (2021).

### 4. Differential expression analysis

The strand-specific read counting of the aligned bam files was performed with the featureCount software included in the Rsubread package (Liao et al., 2019). The resulted normalized counts (median of ratios) (Anders & Huber, 2010) were used for differential expression analysis with DEseq2 (Love et al., 2014). Differentially expressed genes (DEGs) were declared as having a Bonferroni’s adjusted *p* value < 0.05. Additional genes not reaching the above-mentioned threshold were analyzed (t-test) and plotted by means of an in-house R script (R Core Team, 2021).

### 5. Real-time quantitative PCR (RT-qPCR) expression analysis

Total RNA from each of the six grape samples described above was used for RT-qPCR. Protocols for cDNA synthesis and RT-qPCR were performed according to Lijavetzky et al. (2008) using a 7500 Real-Time PCR System (Applied Biosystems, Life Technologies). Quantification of miRNAs was performed by means of stem-loop RT-qPCR (Varkonyi-Gasic et al., 2007). Non-template controls were included for each primer pair, and each RT-qPCR reaction was completed in triplicate. Expression data were normalized against the grape *VviUBI* (*Vitvi16g01364*) gene. Relative quantification was performed by means of the ΔΔCt method using the ‘pcr’ R package (Ahmed & Kim, 2018). Gene-specific primers were designed using the Primer Blast web tool (Ye et al., 2012) and the sequences are described in Supplementary Table 1.

## Results

### 1. RNA-seq data mining, identification and annotation of phase transition-related genes

Given the established role of various miRNA families in the genetic regulation of *Arabidopsis* vegetative phase transition (Poethig, 2010; Wu & Poethig, 2006; Xu, et al., 2016), a fundamental prerequisite for conducting a comprehensive RNA-seq transcriptomic analysis of the vegetative phase transition in grapevines is to obtain the corresponding annotation for these miRNA families’ sequences. As this info was not present in the original annotation file corresponding to the PN40024.v4 genome assembly used in the present work (*Vitis_vinifera - Ensembl Genomes 59*, n.d.; Yates et al., 2022), we enriched that annotation by assembling the unannotated transcripts generated in the present experiment (see Materials and methods “Library construction and RNA sequencing”) and mainly by adding an miRNAs annotation file available at The miRBase Sequence Database (Release 22.1; https://www.mirbase.org/download/vvi.gff3). Due to their relationship with plant phase transition, we focus on three families of miRNAs sequences: miR156, miR172, and miR159. In Supplementary Table 2 we described the 15 miRNAs added to the PN40024.v4 genome annotation, nine miR156, four miR172, and two miR159. Just one miRNA sequence (i.e., vvi-MIR156d) was already present in the original annotation (Vitvi11g04173), while the annotation of vvi-MIR156g was original from the present experiment. Additionally, we also performed the re-annotation of the putative SQUAMOSA PROMOTER BINDING PROTEIN-LIKE (SPL) transcription factors identified in the PN40024.v4 genome. Using the information reported by Hou et al. (2013) and performing protein homology searches of the sequences of the *Arabidopsis SPL* genes against the NCBI (Wheeler et al., 2006) and Ensembl Plants (Yates et al., 2022) databases, we came-out with a final set of 16 putative grapevine *SPL* genes (Supplementary Table 3).

To evaluate the intra- and intergroup sample variability we performed the analysis of the global gene expression of the six evaluated libraries (three replicates for each developmental stage). The Principal Component Analysis (PCA) plot (Figure 1A) showed that the variability between groups was much greater than that observed within the groups, where PC1 was clearly the main source of variation (73.8%), while the variation within replicates (PC2) was just 12.6% (Figure 1A). This variation was slightly larger in juvenile replicas, likely related to the fact that these samples were not identified by a common marker like the appearance of the tendril primordia used for the adult replicas.

**Figure 1.**
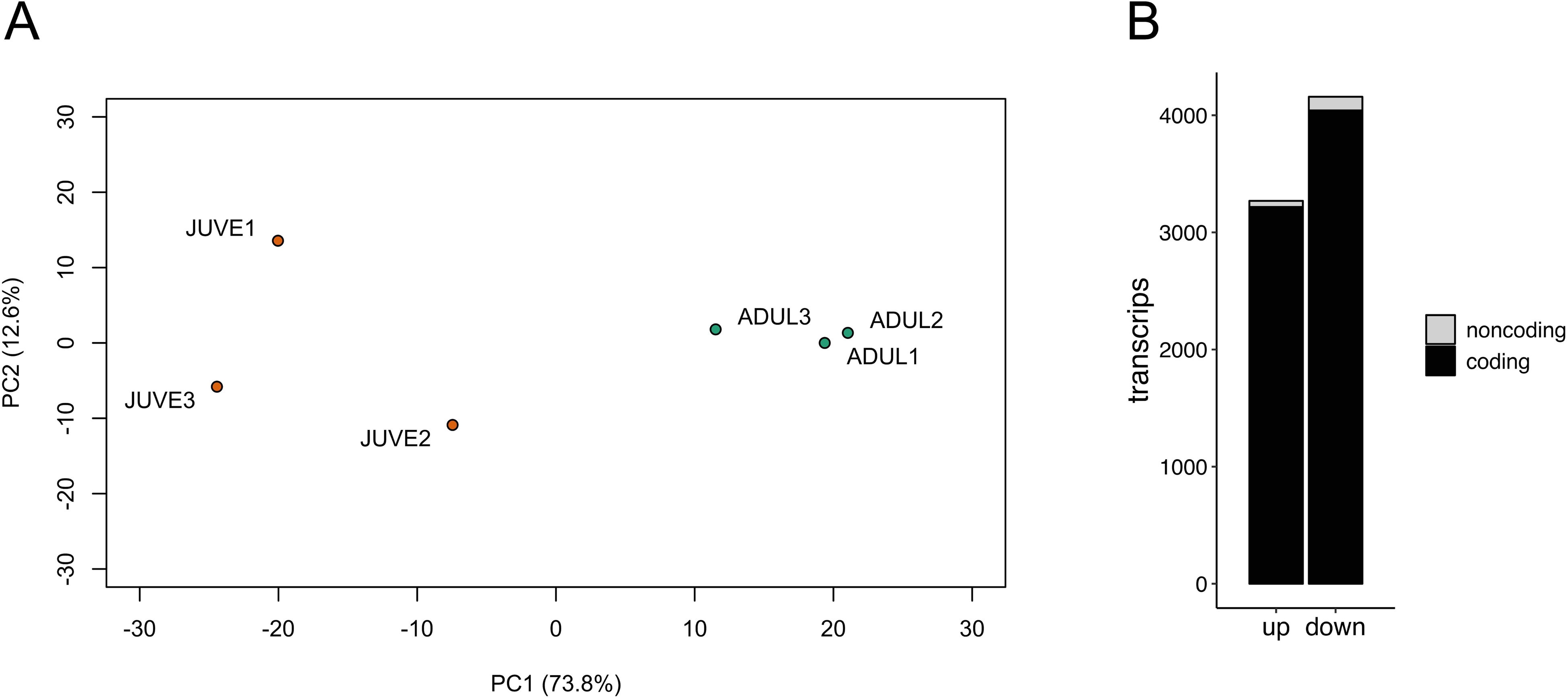
Expression and annotation of phase transition-related genes. (A) PCA analysis of the global gene expression of the six evaluated libraries (three replicates for each developmental stage). (B) Differentially expressed genes (up- and down-regulated) between juvenile and adult plants (Bonferroni’s adjusted *p* value < 0.01) distributed by coding and noncoding transcripts.

We then assessed the variation in gene expression between juvenile and adult plants by means of an RNA-seq analysis. A total of 7434 genes were differentially expressed between juvenile and adult PN40024_SP plantlets (Bonferroni’s adjusted *p* value < 0.05; Figure 1B). From those, 7099 corresponded to sequences annotated into the PN40024.v4 genome (Velt et al., 2023), while 335 sequences corresponded to newly assembled and annotated transcripts (see above). New transcripts are identified with the “MSTRG.XXXX” prefix (Supplementary Table 4). Within the DEGs, we found 3274 upregulated and 4160 downregulated transcripts. In turn, the upregulated transcripts were distributed into 3217 coding and 57 noncoding transcripts, while the downregulated ones were divided into 4160 and 119 coding and noncoding transcripts, respectively (Figure 1B). All information concerning the differentially expressed analysis and gene annotation is detailed in Supplementary Table 4.

### 2. Expression of the phase transition regulatory modules miR156-SPLs and miR172-AP2 in grapevine

As expected from previous reports in different herbaceous plant species (Wang et al., 2011; Wu et al., 2009; Wu & Poethig, 2006; Xu, Hu, Smith, et al., 2016, Jeyakumar et al., 2020), miR156 was highly abundant in grapevine seedlings and decreases during the juvenile-to-adult transition. We found a clear repression of four out of nine miR156 sequences (Supplementary Table 2), showing fold-changes (FC) β 5.0. The three miR156 sequences displayed in Figure 2A (i.e., *miR156g*, *miR156d*, and *miR156f*) were differentially expressed in the RNA-seq analysis, while *miR156b* displayed also a clear repression but the values didn’t reach the significance threshold (Supplementary Figure 2). Concerning the putative role of the miR159-MYB module in the regulation of *miR156* (Guo et al., 2017) in the present work, no differential expression was detected for *miR159* or for *Vitvi13g01266*, the putative *MYB33* grapevine orthologue (Supplementary Figure 2), suggesting that grapevine may display a different mechanism of miR156 repression and initiation of phase transition. In this sense, we detected the significant induction of *VvHXK1* (Figure 3), associated in Arabidopsis with the glucose-induced repression of miR156 and the consequent promotion of vegetative phase change (Yang et al., 2013).

**Figure 2.**
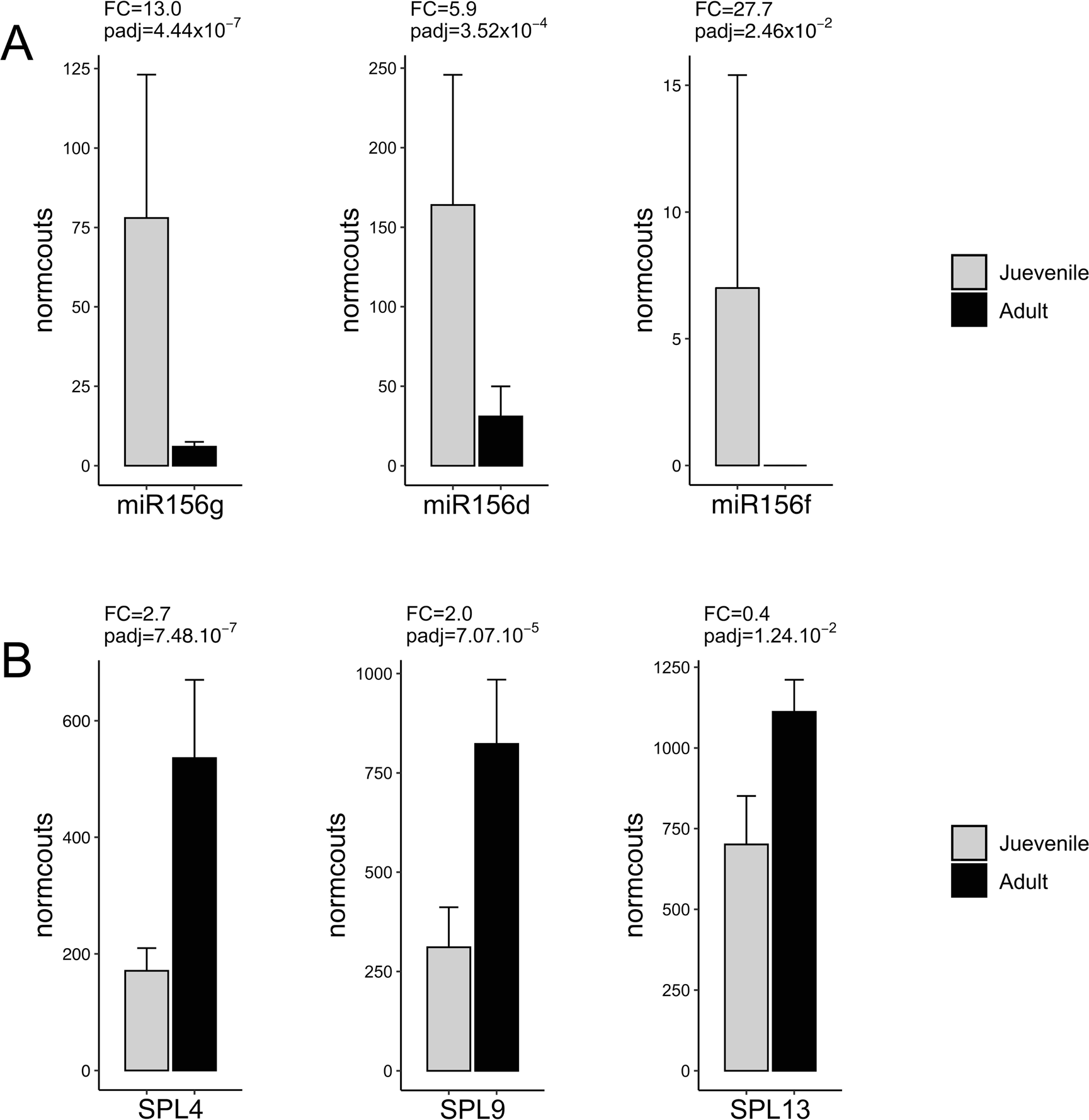
Differential expression (FC= fold-change) between adult and juvenile plants for three miRNA156 (A) and three SPL (B) genes. Data are means of (normcounts = log10 of normalized counts) ± SD of three biological replicates. Adjustment of *p*-value (padj) using the False Discovery Rate (FDR) statistical method.

**Figure 3.**
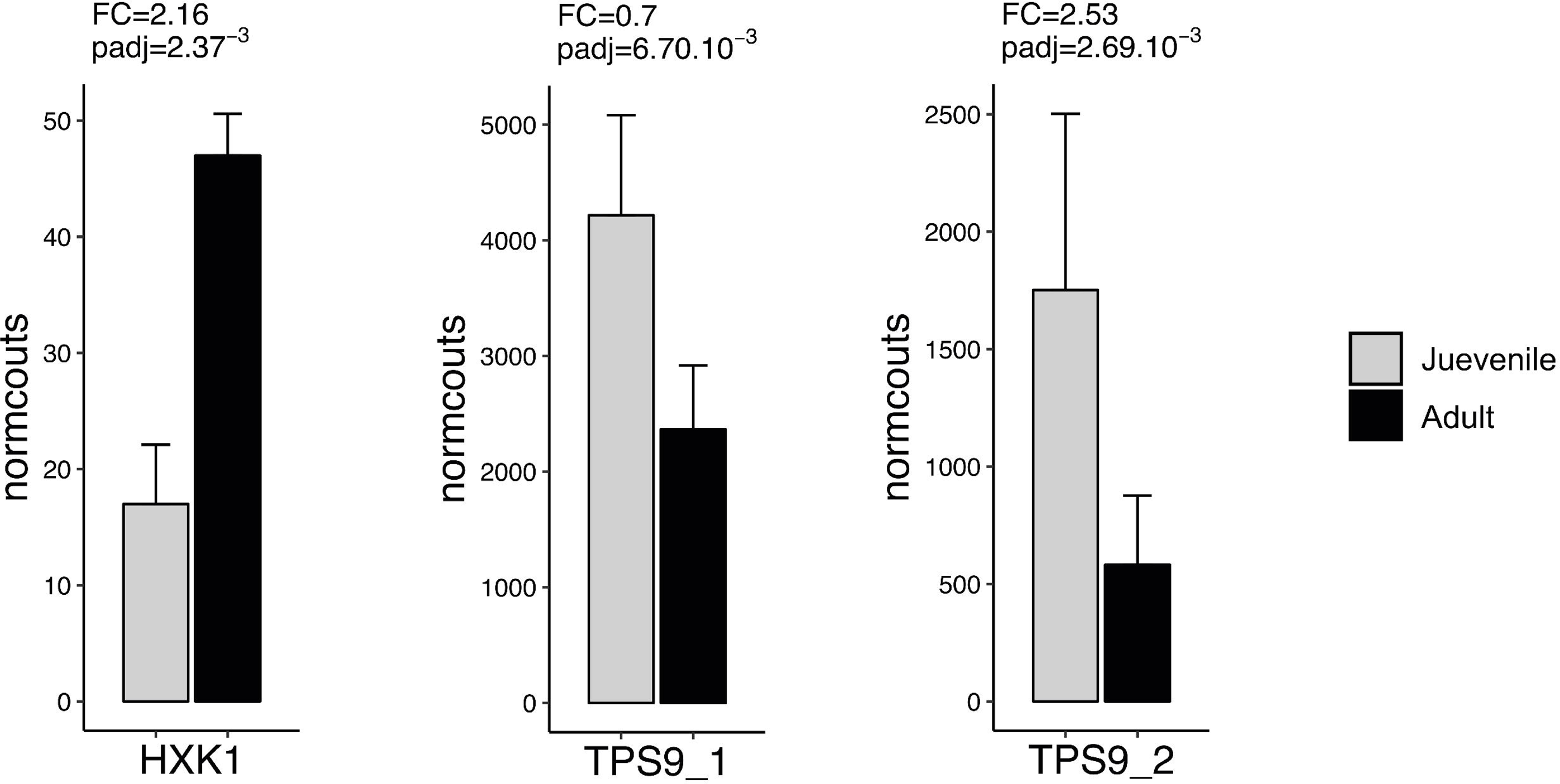
Differential expression (FC= fold-change) between adult and juvenile plants for three sugar signaling-related genes. Data are means of (normcounts = l og10 of normalized counts) ± SD of three biological replicates. Adjustment of *p*-value (padj) using the False Discovery Rate (FDR) statistical method.

It is well documented in *Arabidopsis* that miR156 is essential for managing the timing of the transition from juvenile to adult by coordinating the expression of different pathways that govern various facets of this process (Wu et al., 2009; Wu & Poethig, 2006; Xu, Hu, Zhao, et al., 2016). The miRNA *miR156* acts by inhibiting the expression of different SPL transcription factors during the juvenile phase, so they can express and fulfil their functions right after the repression of miR156 in the adult phase (Wu et al., 2009; Xu, Hu, Zhao, et al., 2016). Accordingly, three *SPL* genes were found induced in adult vegetative grapevines. As shown in Figure 2B, the putative grapevine orthologues of *SPL4*, *SPL9* and *SPL13* were found differentially expressed and showing and opposite behaviour respect to *miR156*. In Arabidopsis, *miR156* suppresses the induction of flowering by inhibiting the expression of *miR172* throughout the repression of *SPL9/10* genes, leading to an increase in the production of AP2-like transcription factors regulated by *miR172*. These transcription factors then suppress the expression of floral activators like *SOC1* (Xu, Hu, Zhao, et al., 2016; Yant et al., 2010). However, despite the significant induction of *SPL9* observed in Figure 2B, no induction was detected for *miR172b* or any of the four *miR172* annotated sequences, while just few counts were detected for *miR172c* (Supplementary Table 2). Accordingly, no repression was observed for *Vitvi06g00360*, the putative grapevine *TOE1* gene (Supplementary Figure 2). It is important to mention that *TOE1* and *TOE2* are the two main AP2-like genes studied in this process (Wu et al., 2009; Xu, Hu, Zhao, et al., 2016), of which only *TOE1* is present in grapevine.

### 3. Expression of genes homologous to Arabidopsis GA flowering induction pathway-genes in the grapevine apex during vegetative phase transition

Despite the facts that we could not detect expression of the AP2-like-miR172 module in the first year of the adult phase and that grapevine plants did not flower during this first year, the floral induction gene *SOC1* showed a slight but significant induction in those adult plants (Figure 4). Moreover, the flowering signal integrator gene *FT*, as well as genes regulating floral meristem identity transition (*AP1*, *LFY*, and *FUL*) showed a consistent and significant expression after the vegetative transition (Figure 4). In addition, as recently reported in *Arabidopsis* (Hu et al., 2023), we detected downregulation of *ZFP21* (Figure 5), the putative grapevine orthologue of *AtZFP8*. This gene, encoding an *Arabidopsis* C2H2 ZINC FINGER PROTEIN (ZFP), release the repression of flower organ homeotic genes after its repression by *AP1* and *LFY*. Interestingly, we also observed the upregulation of *AGL6* and *SEP2* genes in adult vegetative plants, even in the absence of flowering induction (Figure 4). On the other hand, *TERMINAL FLOWER1* (*TFL1*) was the fourth most upregulated gene (Figure 4 and Supplementary Table 4, in agreement with its role of maintaining shoot apical meristem indeterminacy (Hanano & Goto, 2011).

**Figure 4.**
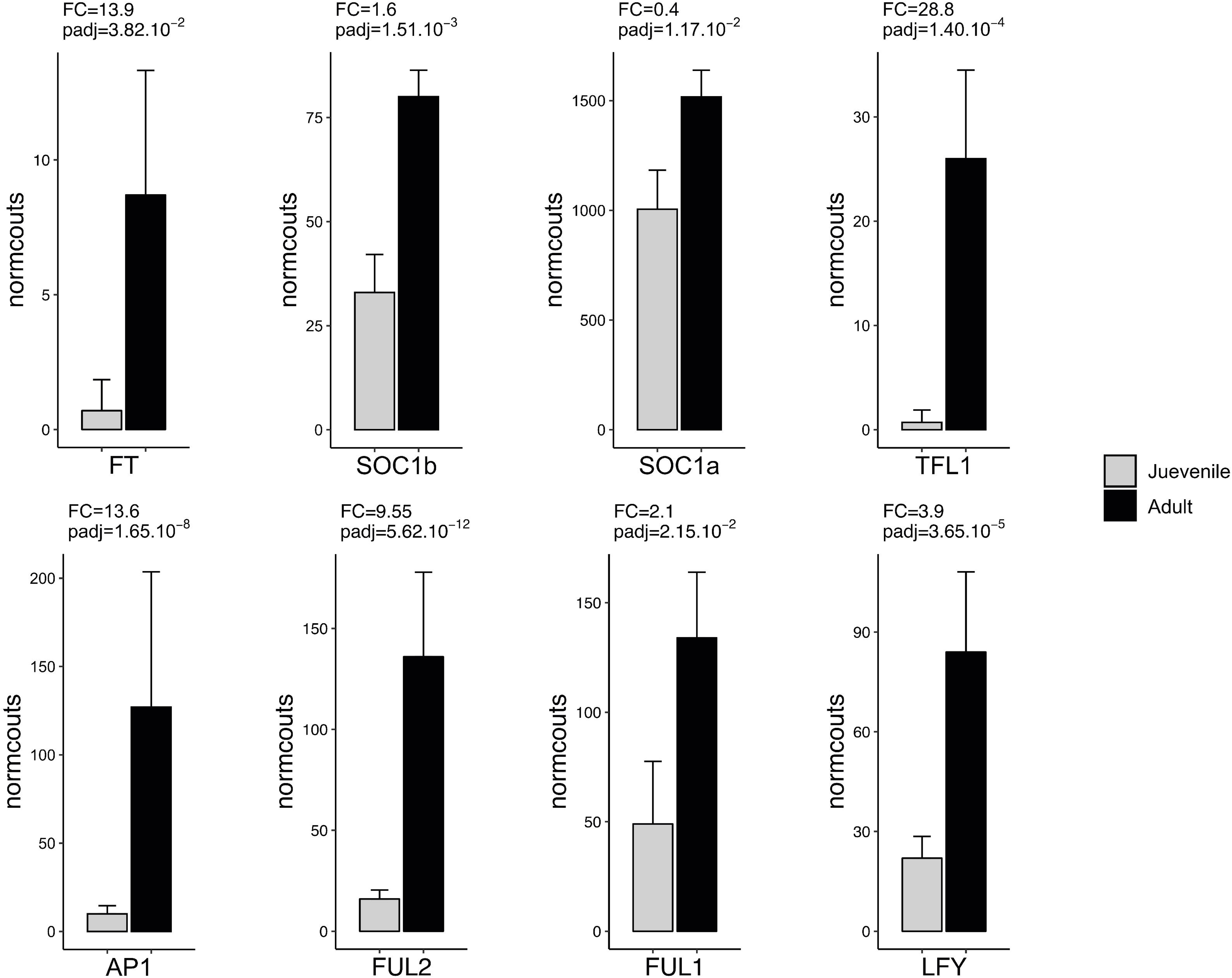
Differential expression (FC= fold-change) between adult and juvenile plants for eight flowering-related genes. Data are means of (normcounts = l og10 of normalized counts) ± SD of three biological replicates. Adjustment of *p*-value (padj) using the False Discovery Rate (FDR) statistical method.

**Figure 5.**
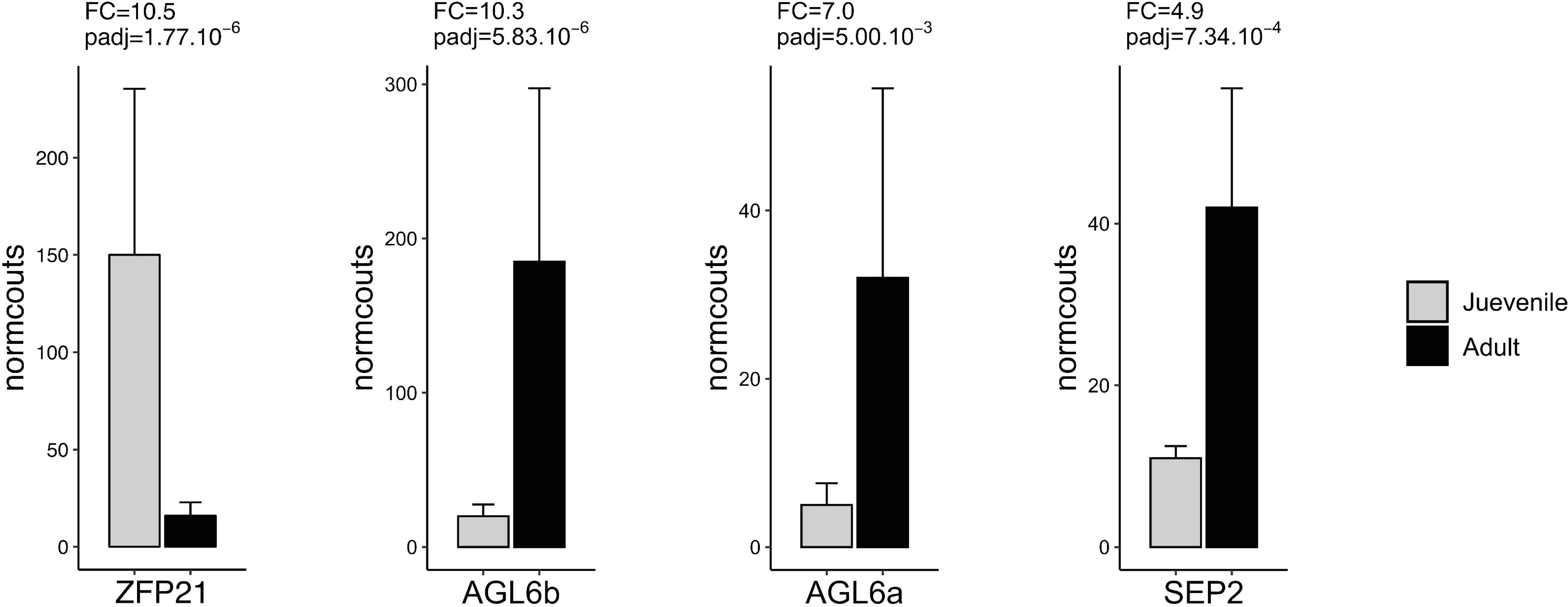
Differential expression (FC= fold-change) between adult and juvenile plants for three class B and C floral homeotic genes and its putative regulator ZFP21. Data are means of (normcounts = log10 of normalized counts) ± SD of three biological replicates. Adjustment of *p*-value (padj) using the False Discovery Rate (FDR) statistical method.

The effects of bioactive gibberellins (GAs) in the control of flower initiation of adult grapevine plants have been previously reported (Boss & Thomas, 2002; Carmona, Cubas, et al., 2007; Carmona et al., 2008; M. Keller, 2020a; Srinivasan & Mullins, 1980). However, the GA involvement in the regulation of juvenile to adult phase transition is unknown. Here, the overall expected effect of the grapevine orthologous of the *Arabidopsis* GA-related genes seems to be a low level of bioactive GAs in the adult vegetative plant and a reduced responsiveness to GAs. As shown in Figure 6, *GA2OX8* was induced, while *GA20OX1* and three positive regulators of GA signalling in *Arabidopsis* (i.e., *SLY1*, *GID1A* and *GID1B*).

**Figure 6.**
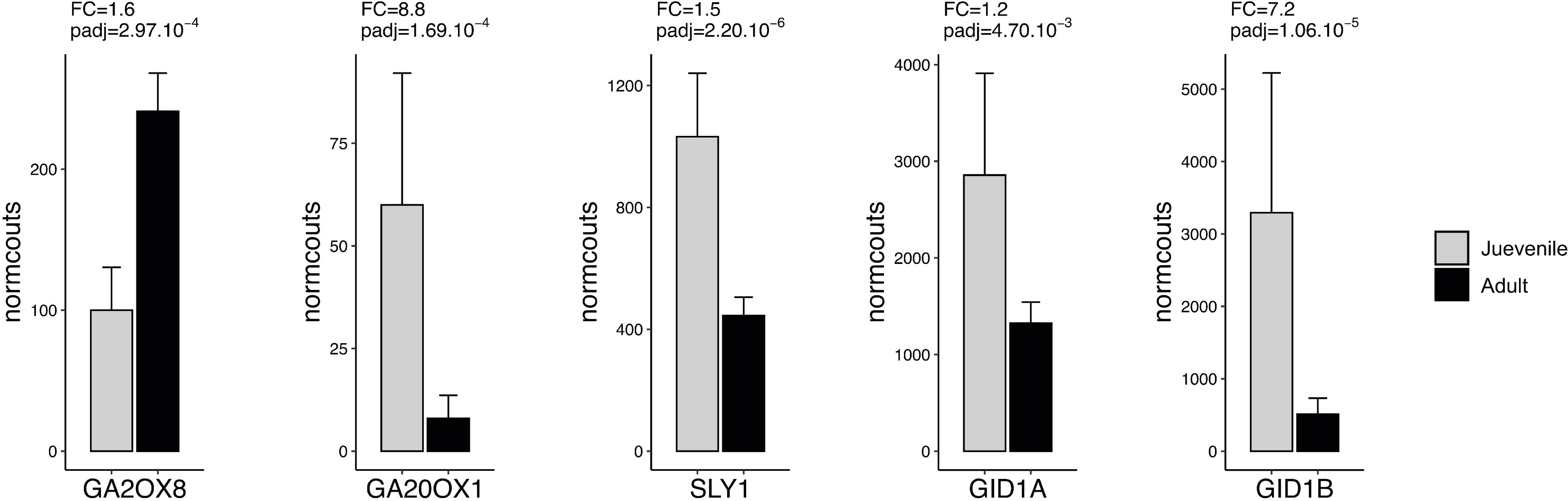
Differential expression (FC= fold-change) between adult and juvenile plants for five genes involved in gibberellins metabolism regulation. Data are means of (normcounts = log10 of normalized counts) ± SD of three biological replicates. Adjustment of *p*-value (padj) using the False Discovery Rate (FDR) statistical method.

### 4. Second year plants

After the initial experiment described above, plants were maintained in the greenhouse one additional year. Then we performed a similar tissue collection as before, harvesting the shoot apex of two sets of plants before and after observing the presence of the first tendril. The RT-qPCR analysis of six selected genes is displayed in Figure 7, where all of them behaved similarly as in the first-year plants. We detected an increased repression of *miR156f* after the phase transition (presence of the first tendril), where again, *miR159c* seems not to be playing a significant role in *miR156c* downregulation (Figure 7). As in the first-year plants, *SPL4* and *SPL9* were significantly induced after *miR156c* repression, but without regulating the expression of *miR172c*. As during the first year, *miR172c* displayed low expression level and no significant induction (Figure 7). Finally, even though these second-year plants also did not show an induction to flowering, the floral meristem identity transition gene *AP1*, also showed a consistent and significant expression after the vegetative transition (Figure 7).

**Figure 7.**
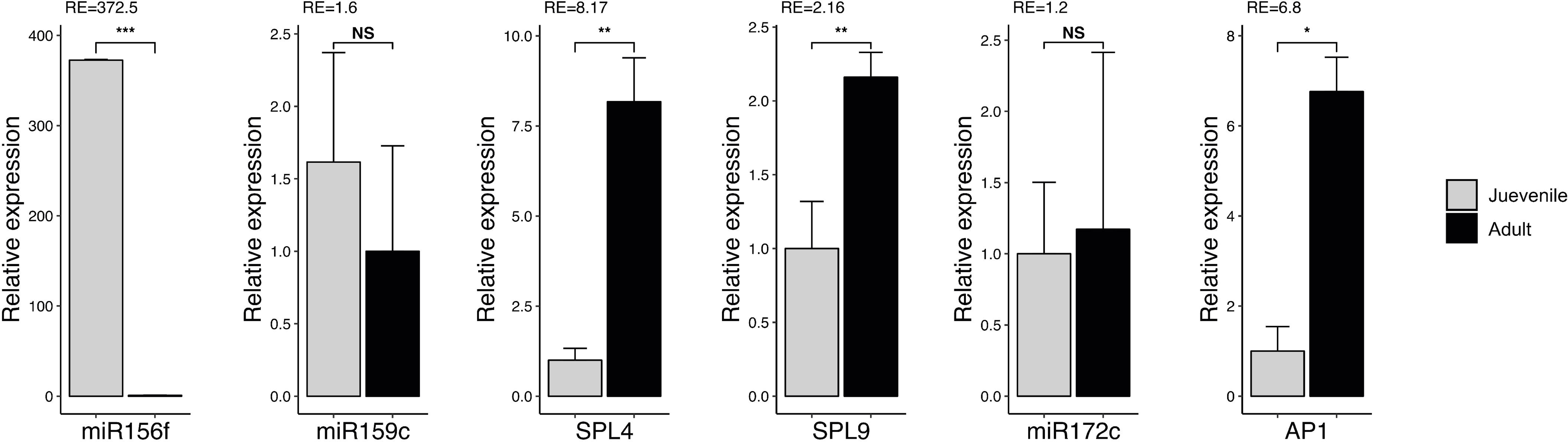
Relative expression (RE) between adult and juvenile plants for six selected genes analyzed by RT-qPCR during the second year. Data are means ± SD of three biological replicates. The *VviUBI* gene (Vitvi16g01364) was used as reference gene. NS not significative; *= *p*<0.05; **= *p*<0.01; ***= *p*<0.001.

## Discussion

The study of phase transition in plants, referred as the progress of the shoot apex through juvenile and adult phases of vegetative development before undergoing a transition to reproductive development is crucial for several biological, breeding, and ecological reasons (Bäurle & Dean, 2006; Poethig, 2010). This process provides an excellent model to study fundamental biological processes such as gene regulation, hormonal signalling, and epigenetic modifications (Lawson & Poethig, 1995a; Xu, Hu, Smith, et al., 2016). Moreover, understanding the molecular mechanisms behind the transition to flowering helps in breeding crops that flower at the right time, maximizing yield in diverse environmental conditions. (He et al., 2020; Kinoshita & Richter, 2020). With changing global climates, knowledge on the biology of phase transition is the basis for the development of plants that are resilient to altered photoperiods, temperatures, and other stresses (Footitt et al., 2020).

The first targets for studying the existence of distinct juvenile and adult phases of vegetative development in plants were tree species, due to their typically prolonged and stable vegetative growing periods. However, the knowledge about the development and molecular regulation of these processes in perennials remains less studied than in annual plants, where the studies in maize (Poethig, 1988) and *Arabidopsis* (Poethig, 2010; Wang et al., 2011) provided the foundation in this field. Still grapevines, presents various additional particularities regarding the biology of phase transition. In addition to the change in the phyllotaxis of the seedlings (from spiral to alternate) and the morphological change from juvenile to adult leaves, the vegetative phase transition is particularly characterised in grapevines by the appearance of tendrils (Carmona, Cubas, et al., 2007; Mullins et al., 1992). Moreover, in grapevines (and most perennial crops), the vegetative transition of the plant from the juvenile phase to the adult phase and the induction of flowering occur at different seasons, even after two or three years of the seed germination and the corresponding seedling development (M. Keller, 2020b; Wang et al., 2011). These facts, together with the very limited existing literature on the biology of the grapevine plant during its first years of development, condition both the study of physiological responses of the plant and their gene regulation under such transition. The knowledge of the molecular basis of phase change in grapevine may be useful to understand the basis of its genetic variation and its possible relationship with the plant’s fertility in different traits, like the number of inflorescences per shoot. Also, to what extent the phase state can affect the expression of reproductive traits.

Performing a whole genome transcriptomic analysis of the vegetative phase transition in grapevine plantlets grown from seeds presents several challenges. Given the high heterozygosity of cultivated grape varieties, the seeds generated in a cluster are all genetically different (M. Keller, 2020a; Martínez-Zapater et al., 2010). This led us to select the PN40024 line, the only quasi-homozygous genotype available in grapevine (Jaillon et al., 2007; Velt et al., 2023), from which we developed a self-cross population for the transcriptomic experiments.

The results of the transcriptomic analysis of juvenile to adult phase transition provided some insights into the complex regulatory networks underlying this developmental shift in grapevine. These results align with previous studies in model herbaceous annual plants such as *Arabidopsis*, rice, maize or tomato, where downregulation of *miR156* and the consequent upregulation of SPL transcription factors were established as key drivers of the juvenile to adult phase transition (Poethig, 2010; Wu et al., 2009; Wu & Poethig, 2006; Xu, Hu, Zhao, et al., 2016). Likewise, the promotion of phase change, based on the repression of miR156 mediated by sugar (Yang et al., 2013), also seems to be conserved in grapevine. However, grapevine plants display features that might be distinctive. Particularly, while Guo et al. (2017) suggested that repression of miR156 by miR159 regulates the timing of the juvenile-to-adult transition in Arabidopsis by repressing MYB33 expression and preventing this R2R3 MYB domain transcription factor from hyperactivating *miR156* expression throughout shoot development, grapevine initiation of phase transition does not seem to be related to a repression of miR156 by miR159 and MYB33. Moreover, the observed lack of miR172 activation in grapevine as miR156 decays, even with significant SPL gene upregulation, underlines a notable difference from the regulatory interactions observed in annual plants. In *Arabidopsis*, miR172 is consistently activated following miR156 repression, facilitating the transition to flowering (Wu et al., 2009; Xu, Hu, Zhao, et al., 2016). However, in the present work, we observe a significant expression of genes like *SOC1*, *FT*, *AP1*, *LFY*, and *FUL* in adult vegetative plants, despite the absence of *miR172* activation. This may suggest that these genes may be primed for alternative developmental stages rather than immediately driving floral induction. This observation is particularly relevant given the fact that grapevine, unlike annual species, does not initiate flowering within its first year, but accommodates its vegetative development to a perennial lifecycle with annual reproductive events. The slight but significant induction of *SOC1, FT* and the strong upregulation of other flowering-related genes in the apex of adult vegetative grapevine plants may thus could serve as a priming mechanism, setting the stage for later reproductive development and/or to establish a default developmental program of on the lateral meristems for the development of tendrils (Carmona, Cubas, et al., 2007; Mullins et al., 1992). In this sense, the extremely high expression of *TFL1*, the fourth more induced DEG in the apex of adult vegetative plants, seems to agree with its putative role in the maintenance of meristem indetermination in grapevine, resulting in a flowering delay as well as in the development of tendrils (Boss et al., 2006; Carmona, Calonje, et al., 2007; Fernandez et al., 2014). In grapevine, the transition from juvenile to adult phase is required early, because the plant needs the tendrils (sterile modified inflorescences) to climb through the riverbank forest to reach the canopy light, but this could take several years (Carmona et al., 2008; M. Keller, 2020b). The adult plant needs light to grow and accumulate reserves and will not flower until it has fulfilled it. In this sense, the low levels of *TPS* could indicate that the plants are not yet ready to begin flowering, in agreement with the reports on the Arabidopsis TPS loss of function plants, showing extremely late flowering (Wahl et al., 2013).

In grapevine (Boss & Thomas, 2002; Carmona, Cubas, et al., 2007; Carmona et al., 2008) and other perennial plants, particularly fruit trees, GAs are generally found to inhibit flowering (Khan et al., 2014; Nishikawa et al., 2009). Under such context, in noninduced adult vegetative grapevine plants, one may expect higher levels of bioactive gibberellins and the plant’s increased sensitivity to respond to GAs. Conversely, here we observe a potential increased degradation of GAs (due to *GA2OX8* up-regulation), a reduced biosynthesis (due to *GA20OX1*) and a putative decreased sensitivity to GAs (due to *SLY1*, *GID1A* and *GID1B* down-regulation), factors more expectable for *Arabidopsis* than for grapevine. A more complete understanding of these phenomena requires references to GA levels and their activity thresholds during the development period under study. An additional experimental limitation is that the apexes may present different levels of GAs in meristems, leaf primordia, and tendril primordia.

In Arabidopsis, flowering is followed by the overexpression of several genes associated with flower induction and development (Blázquez et al., 2001). On the other hand, in grapevine adult apex we found a significant expression of most of the corresponding orthologous genes, but at lower levels that those found in inflorescence meristems (Díaz-Riquelme et al., 2009, 2012, 2014), and not inducing the development of inflorescences and flowers but rather of primordia and tendrils. Even though tendrils are considered organs analogous to inflorescences, supported by their shared developmental origins, functional roles, and regulatory mechanisms (Boss & Thomas, 2002; Calonje et al., 2004; Carmona, Cubas, et al., 2007; Carmona et al., 2008; Srinivasan & Mullins, 1980), it would seem that there are some missing factor(s) that could allow floral induction during the early years of development. In the present work, two main singularities are characterizing plant development and transition. First, the absence of miR172 expression, a key regulator of flowering induction in adult vegetative plants (Khan et al., 2014; Wu et al., 2009), and secondly, the downregulation of *TPS*, highlighting the putative role of carbohydrates-mediated signalling in this step. To what extent these two phenomena are cause and/or effect of the absence of floral induction presumably mediated by other different factors is a question still to be resolved.

## Acknowledgements

This work was supported by European Commission’s Horizon 2020 Marie Sklodowska-Curie-Research and Innovation Staff Exchange programme “(vWISE) Vine and Wine Innovation through Scientific Exchange”, Agencia Nacional de Promoción Científica y Tecnológica (ANPCyT): PICT2018-02381 and PICT-2021-00039; Spanish Agencia Estatal de Investigación PID2020-120183RB-I00. We are grateful to Pablo Carbonell-Bejerano for his helpful comments and discussion, Servicio de Recursos Vegetales (ICVV) for maintenance of the plants and Miguel Angulo for technical assistance.

