## Supplementary figures and images for "Transcriptomic regulation of juvenile-to-adult vegetative phase transition in grapevine"

### Supplementary Figure 1

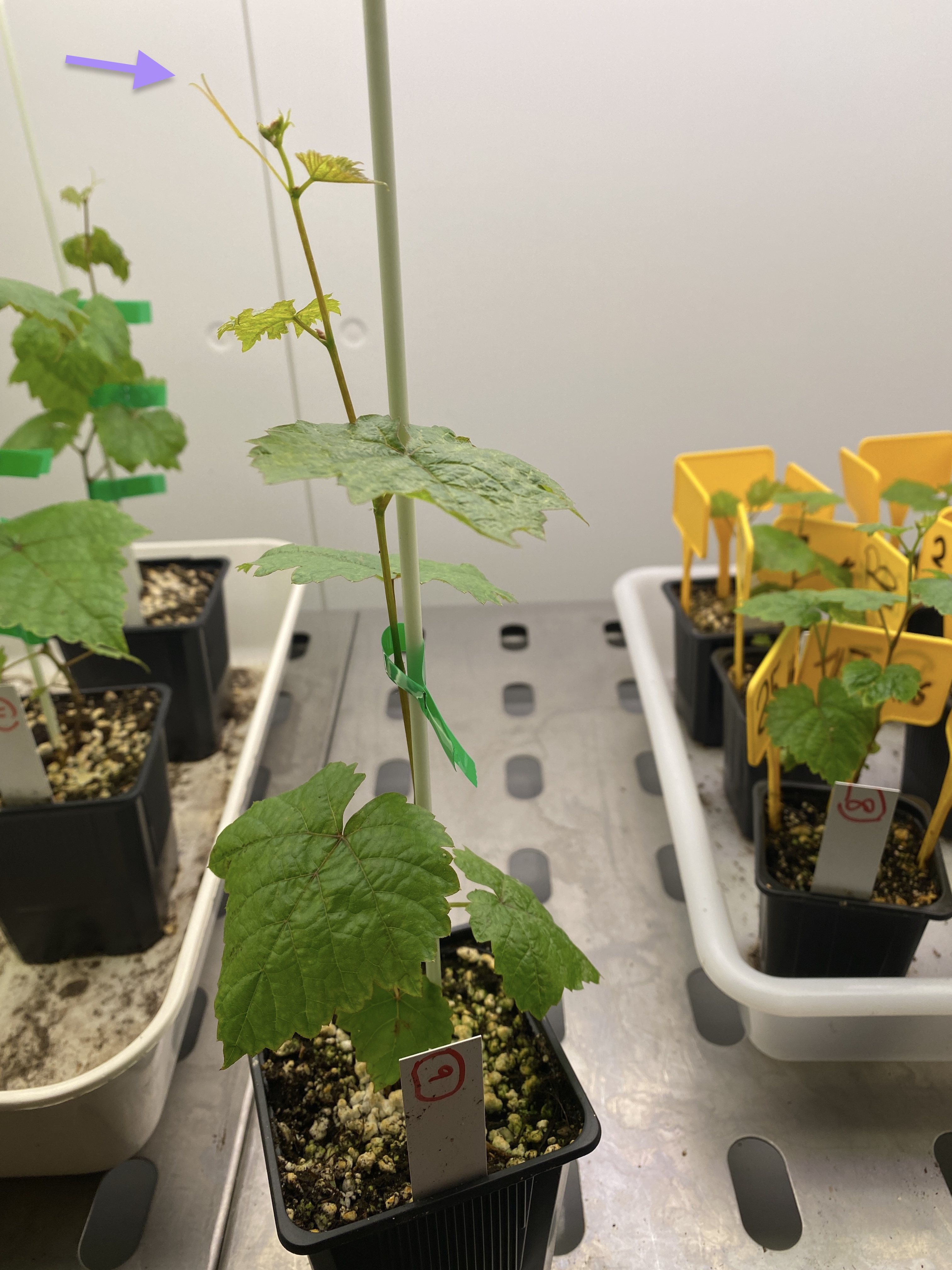

### Supplementary Figure 2

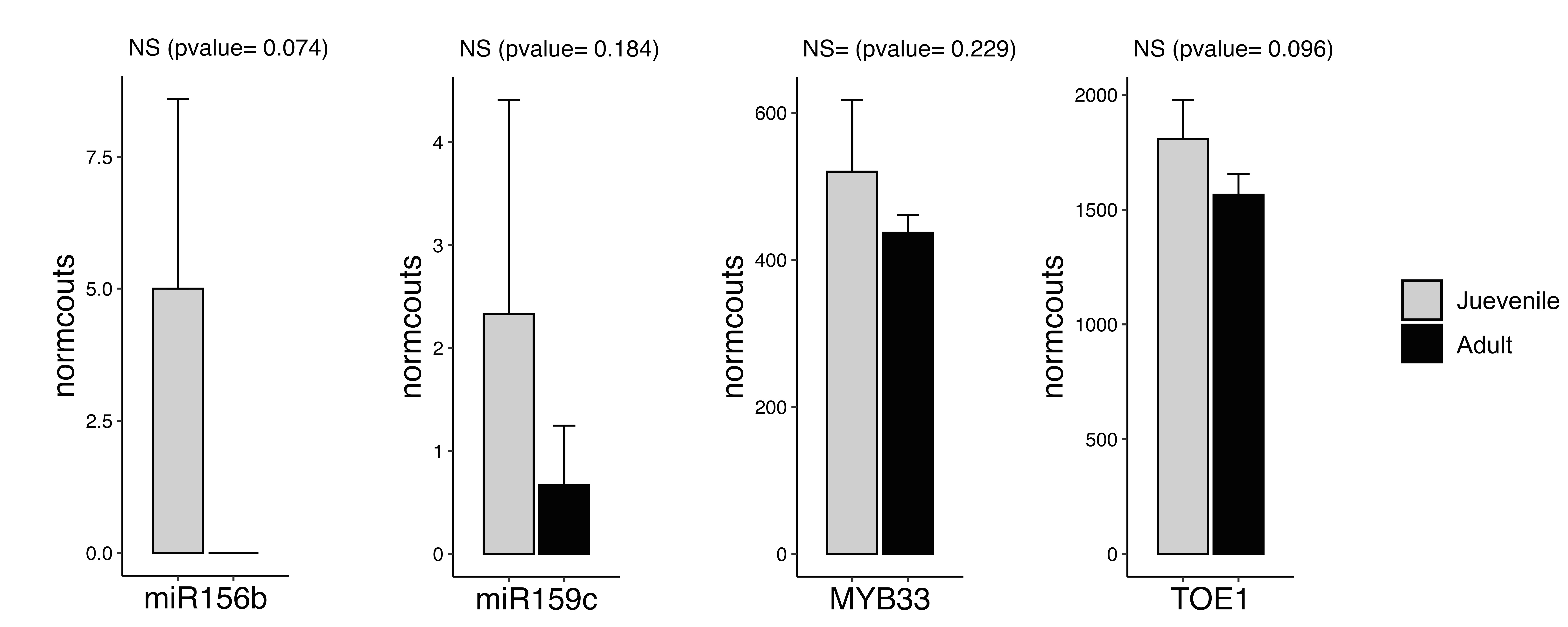
